# A Kinematic Deviation Index (KDI) for Evaluation of Forelimb Function in Rodents

**DOI:** 10.1101/2024.09.26.615237

**Authors:** Abel Torres-Espin, Amanda Bernstein, Marwa Soliman, Juan Sebastián Jara, Yunuen Moreno-López, Edmund Hollis

## Abstract

Rodent models are widely used to study neurological conditions and assess forelimb movement to measure function performance, deficit, recovery and treatment effectiveness. Traditional assessment methods based on endpoints such as whether the task is accomplished, while easy to implement, provide limited information on movement patterns important to assess different functional strategies. On the other side, detailed kinematic analysis provides granular information on the movement patterns but is difficult to compare across laboratories, and may not translate to clinical metrics of upper limb function. To address these limitations, we developed and validated a kinematic deviation index (KDI) for rodents that mimics current trends in clinical research. The KDI is a unitless summary score that quantifies the difference between an animal movement during a task and its optimal performance derived from spatiotemporal marker sequences without pre-specifying movements. We demonstrate the utility of KDI in assessing reaching and grasping in mice and validate its discrimination between trial endpoints in healthy animals. Furthermore, we show KDI sensitivity to interventions, including acute and chronic spinal cord injury and optogenetic disruption of sensorimotor circuits. The KDI provides a comprehensive measure of motor function that bridges the gap between detailed kinematic analysis and simple success/failure metrics, offering a valuable tool for assessing recovery and compensation in rodent models of neurological disorders.

## Introduction

Rodents are often used to model human neurological conditions such as spinal cord injury (SCI), stroke, multiple sclerosis, and Parkinson’s disease, as well as musculoskeletal disorders of bone, joints and muscles. Forelimb movement in these models is assessed using specific motor tasks like reaching for a food pellet as a primary outcome measure reflecting their level of function and evaluating disease progression and treatment effectiveness. These tasks are time-consuming, limiting the amount of detailed information obtained from the summary metrics commonly reported, such as the number of times an animal successfully reaches, grasps, and retrieves a food pellet in a pre-set number of presented pellets or time ^1–4^. The advantage of these summary measures is that they are easy and fast to calculate and do not require specialized equipment. Their drawback is that they only provide information on the task result (e.g., percentage of success) but do not provide qualitative or quantitative measures of the movement patterns that produced the specific result.

Distinguishing between patterns of movement during forelimb tasks is important since different sequences could produce the same result (e.g., reaching a pellet) but provide very different information on the mechanisms underlying the measured function. A common question when using these forelimb tasks as primary outcomes is whether the results of the task are due to recovery of the original movement pattern or the generation of a new one, referred to as compensation^2,5–7^. One simple way to evaluate compensation is by preventing the compensatory movement. For example, rats often develop a scooping mechanism to obtain pellets in the single pellet reaching and grasping task by dragging them on the pedestal, which a gap between the pedestal and the chamber can prevent^8^. Another, more informative, way to understand the nature of the functional outcome is to analyze the sequence of movements during these tasks. The stereotypic movement can be divided into a sequence of well-defined moves (e.g., lifting the paw, reaching) that can be quantified individually ^9,10^ or the full temporal pattern and kinematics can be obtained using motion-tracking systems ^9,11–13^. This requires high-speed video recording and specialized equipment, with complex offline analysis, that limits its widespread adoption. With the advent of machine learning computer vision systems, markerless motion tracking has reduced some of these limitations ^14–17^. However, these systems provide vast amounts of highly granular data (i.e., frame-by-frame position of several markers) that requires analysis using specialized methods and experience. In addition, it is not clear whether the conclusions driven by that highly granular and detailed analysis are transferable to the clinical setting.

In the clinical setting, motion scales and instruments such as the Gross Motor Function Classification Scale^18,19^ or the Fugl-Meyer Assessment^20–22^ are common methods for evaluating movement dysfunction and identifying different levels of disability. They rely on an observant qualifying whether a subject can make a set of movements, which requires well-trained individuals to conduct the tests. Different observant-based methods may result in variable results, with inter-rater reliability varying across methods ^23–25^. An alternative method is to use quantitative indicators derived from overall movement and contrast these indicators against measurements from a healthy reference population. For example, the Gate Deviation Index^26–29^ assesses the dysfunctional gait in different neurological conditions. In upper limb function, Jurkojc et al.^30^ described the creation of a standard deviation of the differential index (SDDI) based on kinematic parameters of objective assessment of upper limb motion pathology. The SDDI determines the standard deviation of the average distance of a movement pattern to a set standard of movements from healthy individuals. A different SDDI is calculated for a time-series trajectory of 9 different movements (e.g., flexion/extension of the elbow joint) and averaged to provide a global summary of deviation from the standard. This requires the selection of a set of pre-determined movements. Movement-agnostic methods have also been described. Recently, Halvorson et al.^31^ described the development of a kinematic deviation index (KDI) for the star excursion balance test in humans in an unsupervised fashion. They embedded the spatiotemporal data of six tracked joints in three dimensions to a three-dimensional space using principal component analysis (PCA) and calculated the deviation from a theoretical subject-specific optimal movement. This effectively captured global function without the need to pre-define a set of specific movements for analysis. Less attention has been given to developing these indices capturing global motor function in animal models of human pathology.

Inspired by previous work on humans^31^ and the need to define metrics in experimental research that mimic the clinic, we describe the development and validation of a KDI in rodents. This metric is a unitless summary score that reflects the multivariate overall movement performance difference to a control condition derived from the spatiotemporal sequence of markers without the need to pre-specify specific movements or features. We illustrate its use in the case of reaching and grasping in mice.

## Material and Methods

### Subjects

For the purpose of metric derivation and validation, we used data from ongoing experiments, whose experimental results will be described elsewhere. All animal work was approved by the Weill Cornell Medicine Institutional Animal Care and Use Committee. C57BL/6J mice (Jackson Laboratory) were housed in disposable plastic cages on a 12-hour light cycle, humidity 39-48%, and temperature of 21.7°C. Experiments were conducted on adult male and female mice. Prior to SCI C57BL/6J mice were food restricted to 80–90% of their free-feeding bodyweight, then tested on the single pellet reach task for the dominant forelimb. Surgical procedures targeted this forelimb.

For optogenetic control experiments, mice underwent cortical transduction of primary motor cortex (M1; A/P 1.5 mm, M/L 1.5. mm from bregma) with 180 nl injection of AAV-retro-hSyn1.Hi.eGFP-Cre (Addgene 105540, 1.0 × 10^13^ GC/ml) and primary sensory cortex (S1; A/P 0.27-0.7 mm, M/L 2.25 mm from bregma, 0.3mm depth) with 270 nl of Cre-dependent AAV1-hSyn1-SIO-stGtACR2-FusionRed (Addgene 105677, 1.0 × 10^13^ GC/ml). A 1.25 mm stainless steel cannula with guide (ThorLabs, CFM22L02) was affixed over S1 using Vetbond cyanoacrylate glue (3M) and C&B-Metabond dental adhesive cement (Parkell). Mice recovered for 1 week following surgery and then were trained on a single pellet reach task, as detailed below. Following training, mice underwent baseline testing sessions with and without IR-triggered optogenetic stimulation. As mice reached through the slot towards the pellet, interruption of an IR beam triggered stimulation with a 473 nm fiber-coupled laser (20 mW). Mice were then subject to a cervical spinal level 7 (C7) dorsal column lesion and allowed to recover for 1 week. Mice underwent 4 weeks of rehabilitation on single pellet reach for 5 days per week, before being tested again with the laser on and off.

For chronic SCI experiments, we used an injury model that specifically targeted ascending sensory circuits carrying tactile, vibration, and proprioception from the affected forelimb as well as descending motor circuits required for skilled motor control, principally the corticospinal and rubrospinal tracts. Following fourteen days of training on the single pellet reaching and grasping task and recording baseline assessment, C57BL/6J mice underwent dorsal over-quadrant section (DoQx) micro-scissor lesion at cervical 7 (C7) spinal level to transect the dorsal columns bilaterally and the dorsolateral spinal cord unilaterally to the dominant forepaw, to a depth of 1 mm from the dorsal surface of the cord. Mice were then returned to the home cage for 12 weeks before single pellet reach performance was tested again.

### Single pellet reach task

Food-restricted mice were placed in an acrylic behavior box with three 7 mm wide slots, positioned on the left, center, and right sides of the front wall. Flavored food pellets (20 mg, Bioserv, F05301) were placed 12 mm away from the inside wall of the box on a platform level with the box floor, a gap prevented mice from sliding pellets directly to their mouths. Mice were trained over 15 sessions (25 trials per session) spread across 3 weeks. For manual quantification of success, only trials with successful pellet targeting were counted, and the success rate was calculated as the percentage of trials with pellet retrieval and eating. High-speed video of the task was recorded on a Basler Ace camera (acA1440-220um) with a 12 mm lens at 327 fps, 720 × 540 px resolution for quantification of forelimb reach trajectories using the markerless pose estimation AI DeepLabCut ^14^.

### Kinematic data collection and processing

Seventeen videos were labelled and used to compose the training data set in DeepLabCut (version 2.2b7). From each video, at least 70 frames were extracted where digits 2-5, and the pellet were labeled. These frames were used to fine-tune the pre-trained network (ResNet-50) to predict features of interest in unseen videos and generate x- and y-coordinates for each label throughout the videos. The raw kinematic data for each separate trial for each animal, consisting of the frame-by-frame horizontal (x) and vertical (y) location for each marker, is pre-processed to eliminate spatial and temporal outliers. For spatial outliers, following biomechanical spatial coherence between markers, the Euclidian distance between markers for each frame was calculated and any point above 92% of the between-marker distance distribution over all frames was removed. For temporal outliers, following temporal coherence, a smoothing regression (Friedman’s Super Smoother) for each raw data trace per marker and coordinate (x and y) was performed using the R forecast package^32^ to reduce noise and detect data points with large residuals reflecting low temporal coherence and substitute them with the expected predicted value from the regression. Any missing value as imputed using the predicted value from the smoothing regression. Given that each trial has a different duration, and therefore a distinct number of frames, each variable was linearly interpolated to 100 sample points, where each sample represented 1% of the temporal sequence of the trial for each marker and coordinate (x and y). The outlier corrected, time normalized and smoothed data was use for deriving the KDI.

### Kinematic Deviation Index (KDI)

For each trial the KDI was derived as follows: the data was organized on an *n x p X*_*i*_ matrix where *i* denotes each individual trial, *n* ={1,…,100} representing the % duration of each recording, and *p* =(1,…,*v*,) where *v* are the variables measured (x and y coordinates per marker, digit span, and angle). A PCA was performed on each trial (*i*) independently. Successful baseline trials were extracted for each trained animal, and a Generalized Procrustes Analysis (GPA) was performed on the loadings of the first 3 PCs across all PCA solutions of the successful baseline trials to rotate all PCAs to a common space (correcting for rotational indeterminacies). The consensus GPA space (average) across successful trials for each animal at baseline forms the subject-specific kinematic performance reference, which is used as an anchor representing the best estimate of pre-perturbation PCA space. We then rotate the loadings of the first 3 PCs of each trial in each animal toward the subject-specific kinematic performance reference using Procrustes rotation. The rotated loadings for each trial are applied to the original *X*_*i*_ by the dot product to obtain rotated object scores, which is a vector of length *n* with the coordinates of the integrated variables over the reference space for each % increment of the trial duration. In that sense, the object scores for each PC for each trial represent a summary of all markers with respect to the kinematic reference. The Euclidian distance for each n sample between the test trial and the kinematic reference on the 3 PCs was calculated, which we refer to as the KD-trajectory, and the sum of these values was used as the KDI per trial:

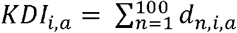 where 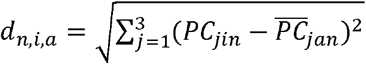 and *a* is the *a*th animal, *j* is the *j* th PC, and 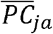 is the subject-specific kinematic performance reference for the *a*th animal and the *j*th PC.

### Statistical analysis

For inference on testing the hypothesis of differences between the levels of a categorical term in KDI we fitted a linear mixed model (LMM) with animal specified as a random effect to account for the repeated measures of several trials per animal, the categorical term as a fixed effect, and KDI as the response variable. In the case of two categorical terms, both marginal effects, as well as the interaction, were considered. An Analysis of Variance Table with Satterthwaite’s method for approximation of degrees of freedom and an F-test were calculated from the LMM. In the case of KD-trajectories (i.e., individual KDI values per frame), an LMM was fitted by specifying a natural cubic spline with 7 degrees of freedom for the frames to consider non-linearity, together with the interaction between the smooth function of frames and a categorical fixed term of interest. Animals were specified as a random effect with random intercept and random slope over frames to account for the individual variance and nested structure of several trials per animal. When indicated, a pairwise contrast was used as a post-doc over the estimated marginal means, and the p-value was adjusted with the Tukey method to compare a family of multiple estimates. The level of significance was set at an alpha of 0.05. Models were fitted in R^33^ using the lme4 package^34^.

## Results

### The kinematic deviation index (KDI)

The use of automatic pose estimation through markerless AI-based methods has greatly augmented the capacity of researchers to obtain high-frequency granular data from tens of markers for kinematic analysis of movement patterns^14,17^. We developed the KDI to produce a behaviorally meaningful metric that integrates the trajectories of distinct markers during single pellet reach in mice into a single metric measuring recovery. The deviation steps can be found in the methods section. Figure 1 visually summarizes the workflow and method. Briefly, after data collection, pre-processing steps are conducted to reduce noise and consider outliers in the marker’s positions derived from the markerless prediction, and to normalize the time-series to a fixed length (supplemental figure). Then, KDI is calculated for each independent trial on the task for each animal, using PCA to embed the spatial position of each marker over time in an overall trajectory (Fig. 1a-c). After the PCA we create a reference of best performance for each animal by aggregating the embedding of successful trials, which the trained animal completed at baseline, to its multidimensional average (Fig. 1d). Our approach assumes that this multidimensional average embedding of successful baseline trials is the optimal sequence of movements that each individual animal has reached in performing the task and serve as a reference to benchmark each subsequent trial. By measuring the deviation of each post-manipulation trial embedding to the subject-specific reference, we can then summarize overall performance trajectory relative to the best expected performance for each animal (Fig. 1e).

**Figure 1.**
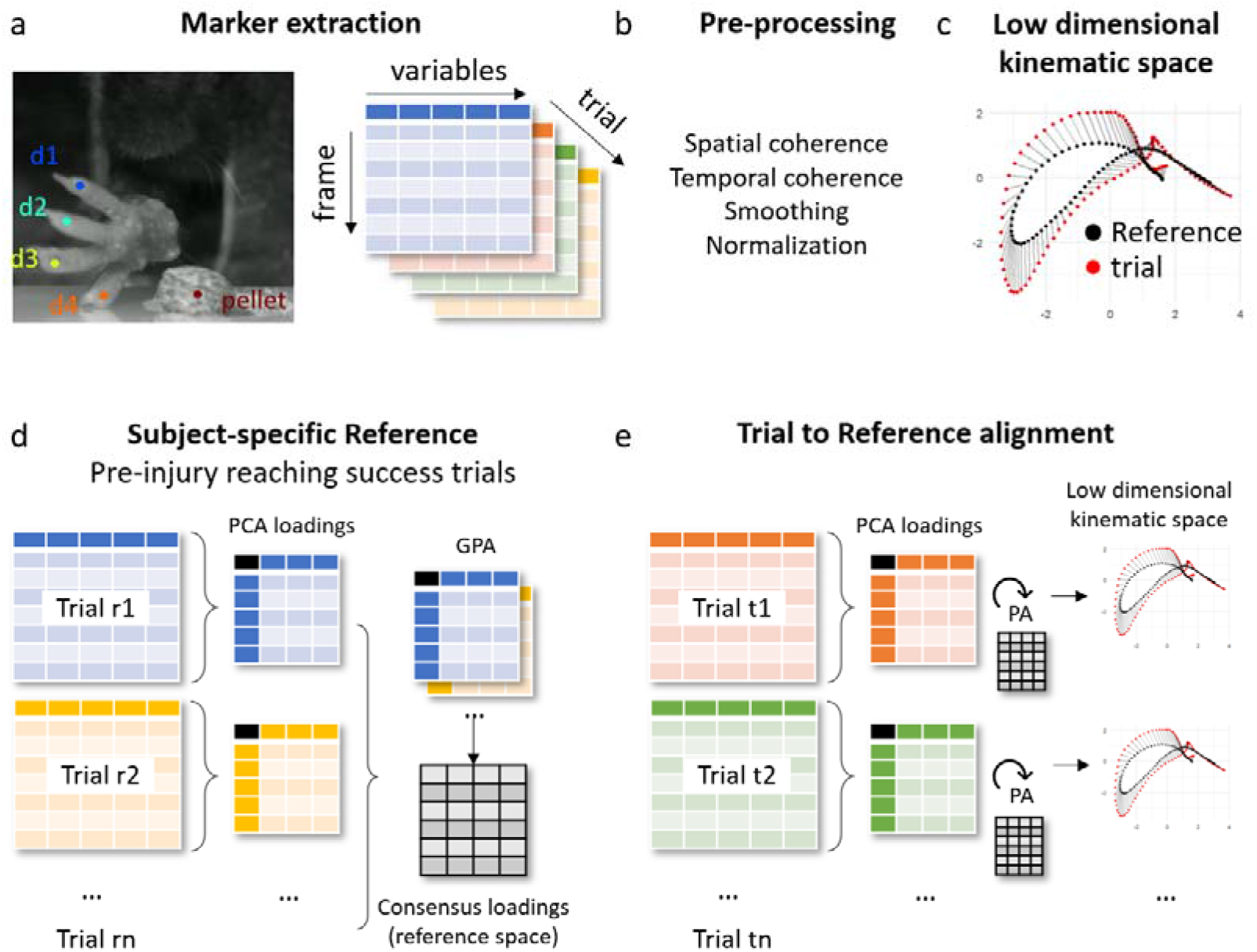
Summary of the workflow for KDI calculation. (a) An illustration of marker extraction and data organization. Each reaching trial constitutes a matrix with frames in rows and marker positions in columns. (b) A sequence of pre-processing steps is conducted to reduce noise and normalize the data in each trial. (c) For each trial, the multidimensional marker data over time is reduced to a low dimensional space capturing the overall trajectory of markers over the trial. (d-e) For each subject, a reference of best performance is calculated from all successful baseline trials and compared to each trial perturbed by experimental manipulation or injury to calculate a global metric of performance.

To quantify the difference between the subject-specific reference and each trial, we calculated two measures, the kinematic deviation trajectory (KD-trajectory), and the KDI. The KD-trajectory measures the difference in location between the reference and a trial for each percent point of the temporal duration of the trial by the Euclidian distance in the embedding space of the PCA. The KDI is the sum of all the differences in the KD-trajectory (see methods) and captures the global deviance between the embedding of a trial to the embedding of the reference. Thus, the lower the KD-trajectory and KDI values, the closer the sequence of movements in a trail to the expected best trained performance of the animal (Fig. 2).

**Figure 2.**
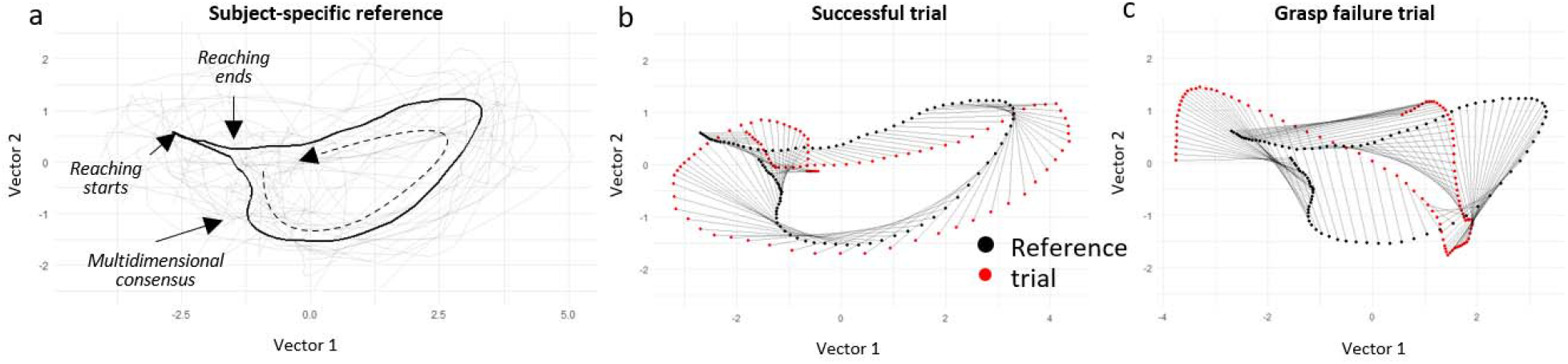
Example of KD-trajectory and KDI. (a) The consensus (black) trajectory of a representative animal over all baseline successful trials (grey), representing the subject-specific reference in the 2D space of the first and second PCs. (b) A successful trial embedded trajectory (red) compared against the reference (black) at each percent point of the trial duration (grey lines between a red and black point). Each grey line represents the deviation of the target trial from the reference in the Euclidian space. The sum of all deviations is the KDI per trial. (c) A grasp fails trail for the same animal, showing the difference in the embedded trajectory between the reference and a failed trial. The dashed black arrow illustrates the directionality of the temporal movement in the embedded space.

### KDI discrimination of single pellet forelimb reach performance

To determine the concurrent validity of the KDI as a metric of overall skilled prehension performance, we compared the KDI values against three different endpoint outcomes during single pellet reach (reach fail: attempt without pellet contact; grasp fail: attempt with successful pellet targeting but not grasping; success: attempt with successful reaching, grasping, and retrieval of the pellet) in mice for two different cohorts of animals trained in the task by two independent investigators (Fig. 3). As expected, KDI values were lowest for the successful trials in both cohorts (Fig. 3a-b). This was true for 90% of the animals, with the 10% remaining having reach failure scores with KDI values within the range of those for successful trials. In both cohorts, KDI values discriminated between trial outcomes, although without reaching statistical significance in the second cohort (pairwise contrast first cohort: p-values < 0.0001 between Success vs Grasping Fail, Success vs Reaching Fail, and Grasping Fail vs Reaching Fail; Second cohort: Success vs Grasping Fail p = 0.064, Success vs Reaching Fail p = 0.098, and Grasping Fail vs Reaching Fail p =0.308). This was also reflected in the KD-trajectories among trials from the same animal (Fig. 3c), providing further details into the differences. For example, for a single animal (Fig. 3c), the KD-trajectory for the successful trials is close to linear, while discrepancies between reach fails and grasp fails from the consensus are more divergent during the first and last quarter of the movement sequence, and again near 50% completion of the movement, when the distance to the reaching target is minimal for successful trials.

**Figure 3.**
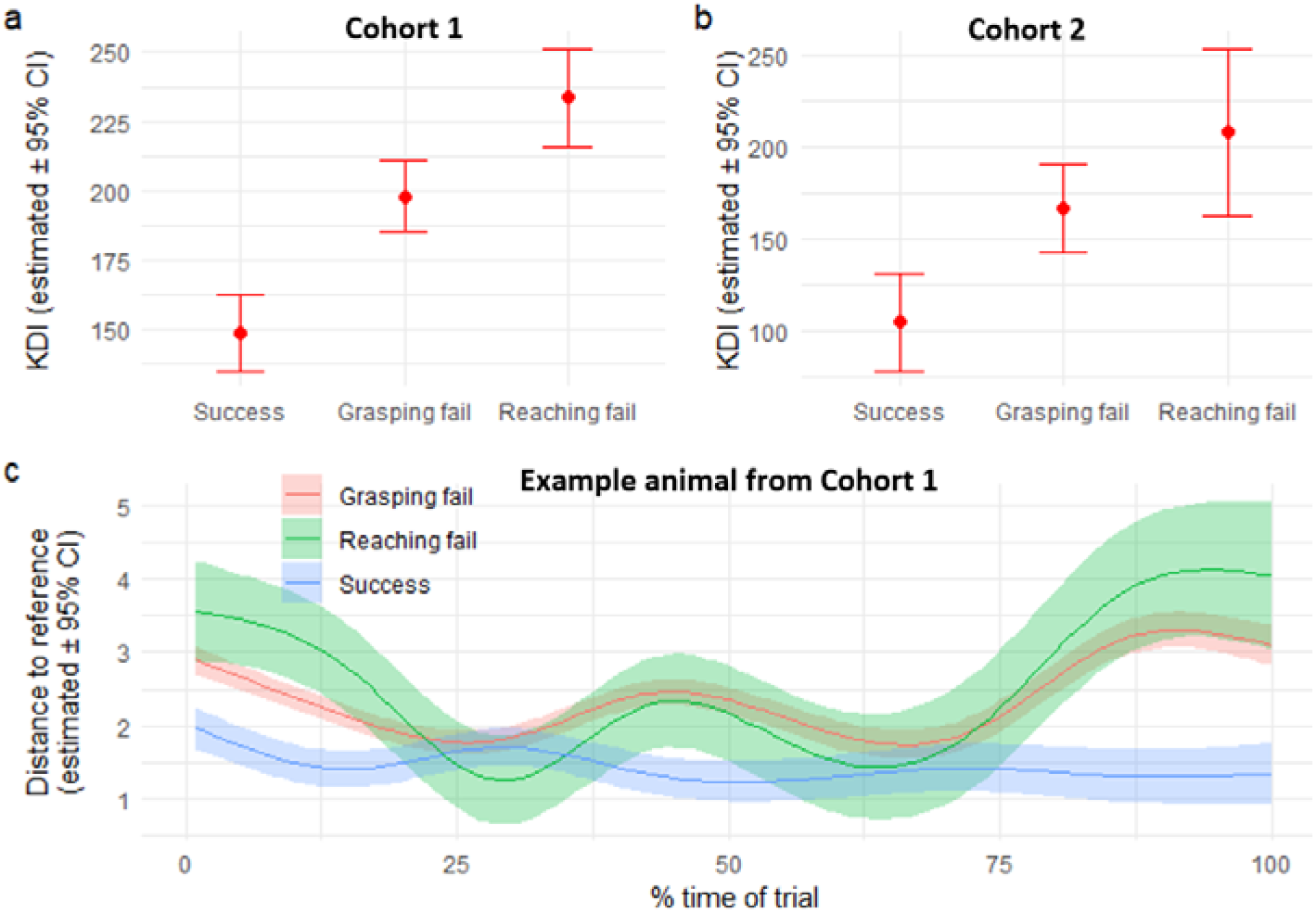
Concurrent validity of KDI. (a) KDI differences between trial endpoints as estimated by a linear mixed model for our first cohort before perturbation (b) KDI differences between trial endpoints for the second cohort before perturbation. (c) A single animal example of the KD-trajectory for different trial endpoints as estimated by an additive mixed model. It provides granularity on the sections of the trial movement sequence where differences with the reference are more prominent for each trial endpoint. Lines and dots represent the estimated values from the analysis models. Error bars and bands represent 95% confidence interval as estimated by the analysis model.

### Sensitivity of KDI and KD-trajectories to intervention

We investigated the sensitivity of the KDI and KD-trajectories to detect changes to two distinct interventions expected to affect forelimb prehension in mice. In our first cohort, we evaluated the effects of chronic spinal cord injury (SCI) on KDI in a mouse model with no expected recovery. Forelimb reach performance was assessed at baseline, prior to SCI, and then again 12 weeks later. The chronic SCI model transected ascending sensory axons and descending corticospinal axons in the dorsal column, as well as the sparse corticospinal component located in the dorsolateral white matter, where descending rubrospinal motor axons were also cut. This injury severely impairs prehension movement, and we observed a significant increase in KDI values (p <0.0001) for all reaches between pre-injury and chronic SCI (Fig. 4a-b).

**Figure 4.**
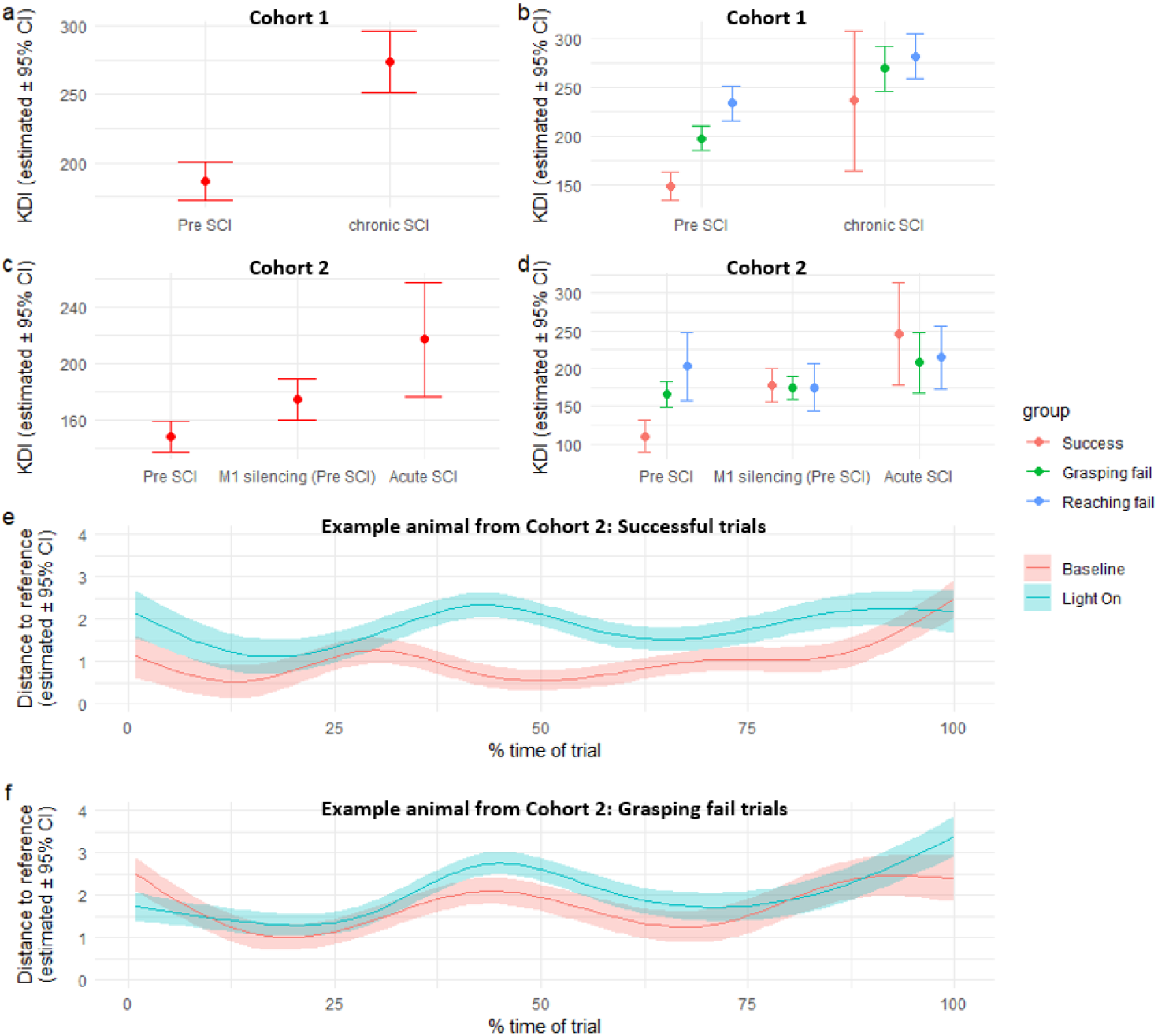
Sensitivity of KDI to experimental perturbation. (a) Differences in KDI per all trials pre-SCI and after chronic SCI and per trial endpoint type (b) as estimated by a linear mixed model for the first cohort. (c) Differences in KDI per all trials pre-SCI, pre-SCI after S1 to M1 silencing (M1 silencing), and acute SCI, and per trial endpoint type (d) for the second cohort. (e) KD-trajectory for a single animal between pre-SCI and when the light is on (M1 silencing pre-SCI) for successful trials, and for (f) grasping fail trials.

In a second cohort of mice, we investigated the effects acute disruption of sensorimotor circuits using optogenetics, in addition to the effects of a more mild, acute SCI. Using an intersectional viral approach, we silenced neurons that project from S1 to M1, carrying afferent information critical to the precise modulation of prehension movements. During silencing, we observed increased KDI for all trials (Fig. 4c; p = 0.03). Notably, the successful trials during silencing presented KDI similar to levels found during reaching and grasping failure (Fig. 4d), suggesting that when sensorimotor circuits are silenced, animals can produce successful trials. Still, those are globally different in movement patterns compared with when the circuits operate uninterrupted. These animals then underwent a dorsal column SCI, that allows for some recovery of success on the single pellet reach task. Testing acutely, 1 week, after SCI showed increased KDI during all trials. Interestingly, the differences between successful trials and those with reach and grasp failures pre-injury are no longer observed after acute (Fig. 4d) or chronic SCI (Fig. 4b).

Comparing the subject-specific KD-trajectories for a single animal before injury and after sensorimotor silencing, provides further insights into differences in movement patterns. For all successful trials (Fig. 4e), silencing the sensorimotor circuits induces an increase in the frame-by-frame distance to the reference. This is especially prominent after the first quarter of the sequence, once the IR sensor triggered laser activation. In contrast, the differences for the trials with failed grasp in the same animal are less apparent (Fig. 4f). This provides further evidence that silencing sensorimotor circuits during a successful trial might disrupt the general trajectory of movement, yet the endpoint of the trial is the same. It also illustrates how KDI and KD-trajectories can be used to capture differences in the global trajectory of movement even among trials with the same endpoint.

Lines and dots represent the estimated values from the analysis models. Error bars and bands represent 95% confidence interval as estimated by the analysis model.

## Discussion

We demonstrate the derivation and validity of a composite summary of the temporal sequence of movements during forelimb prehension in mice using a machine-learning workflow in a similar manner to the proposed metrics in humans. The KDI discriminates between functional endpoints of the single pellet reach task and is sensitive to forelimb functional deficits induced by experimental manipulation, such as injury and targeted disruption of sensorimotor circuits. Moreover, kinematic deviation trajectories provide insight into which movement sequences are disrupted.

Recent advances in automating functional assessment in animals have greatly improved the capacity to conduct more complex experiments with larger sample sizes, increased numbers of experimental groups, and the study of more naturalistic behaviors than previously was possible^35^. Automation can reduce the burden on researchers, leading to increased experimental output, greater throughput per experiment, longer training epochs, and more frequent assessments^1,36,37^. With increased experimental output and the need for accurate, granular behavioral analysis of specific components of sequential movement patterns, high-speed video recording of movements has become essential. In turn, this has created new challenges in robustly analyzing the vast amounts of data collected. Markerless motion pose estimation involves tracking and analyzing the movement of animals without the need for the placement of physical markers. DeepLabCut, one of the most popular tools in fields as diverse as neuroscience, ethology, and biomechanics, leverages deep learning techniques, specifically convolutional neural networks (CNNs), to accurately detect and track key body parts from video data^14^. These tools have grown in popularity to assess all kinds of animal behaviors from video recordings, eliminating the need to instrument the subjects and creating a new field of computational ethology. The use of markerless AI-based methods has greatly augmented the capacity of researchers to obtain high-frequency granular data from tens of markers for kinematic analysis of movement^12,14^. Although this has increased the number of laboratories performing detailed kinematic analysis of movement, it has also increased the challenges for generalizability and reproducibility. Analyzing the large amounts of data generated requires specialized knowledge in kinematic analysis, as well as critical data management and statistics. The lack of a clear framework to synthesize the deluge of data obtained from markerless methods into meaningful metrics of functional deficit and recovery is evident by the various approaches that distinct research groups have used to analyze such data^12,15,38,39^. We developed the KDI to simplify this process and produce a behaviorally meaningful metric that integrates the trajectories of distinct markers during single pellet reach in mice into a single metric measuring function and recovery contrasted to a healthy condition analogous to those used in humans^27,28,30,40^.

Developing methods for quantifying behavior in animals that mimic those used in clinical settings is important for translation, as the mismatch in endpoints between pre-clinical and clinical research contributes to reduced translation^41–43^. The stereotypical movement of reaching and grasping between rodents, non-human primates, and humans is well-conserved^44,45^. Thus, skilled forelimb reaches in rodents are often used as a behavioral output and rehabilitative training in models of human neuropathology, such as SCI, traumatic brain injury, and stroke^2,8,11,13,36,37,46–48^. As it happens with scales and scores in clinical settings, the quantification of movement deficits in animal models with a single endpoint (e.g., pass or fail), or a set of discrete notations of specific movements (e.g., good, partially good, or bad pronation of the hand), is limited by their lack of granularity and subjectivity. A more objective measure is the extraction of quantitative measures from kinematic analysis, such as movement trajectory length, movement duration, mean velocity, peak velocity, and max paw angle, among others^11,15^. One potential limitation of these is the reduction of the trajectory to summary statistics that might not reflect the sequence of movements. Nonetheless, these quantitative measures have been shown to be sensitive to motor strategies after neurological injury^15^, and can serve as general descriptors of the movement characteristics. An alternative approach is to consider the raw information provided by the spatiotemporal trajectories of joints and other landmarks. For example, the MoSeq^49^ approach captures the pose dynamics of animals as they freely move in a variety of experimental contexts by learning independent representations of behavior patterns from the recordings themselves rather than pre-defining what those patterns are. The advantage of this approach is that a set of metrics is not pre-defined, and the totality of the movement data is used in a more data-driven framework, learning important movement subunits that together describe the sequence. Our development of the KDI follows a similar idea in the sense of condensing the spatiotemporal information of all markers into a low-dimensional representation that can be then contrasted against a healthy reference rather than using pre-defined metrics. This parallels the ongoing developments of unsupervised representation learning approaches for evaluating movements aiming to synthesize a summary metric of movement performance^31^.

A major strength of our approach is the integration of the spatiotemporal sequence of several markers into a single metric that discriminates between performance, and it is sensitive to experimental manipulation. The derivation of the KDI can be extended to any set of markers, and the framework we described can be adapted to other tasks using the same function such as the pasta matrix or string-pulling tasks^50,51^, as well as other functions such as locomotion. In addition, these metrics can be implemented in other species such as rats, non-human primates, and humans. Furthermore, our approach is agnostic to the method used for extracting the marker locations and the number of recorded dimensions. Therefore, KDI is applicable to data obtained by diverse instruments and settings, whether through markerless motion detection or infrared motion tracking ^11,14,15^.

One important consideration is the fact that we derive the KDI as a difference to the expected successful trial in healthy conditions per individual. This captures the individual differences in the preferences on performing the task, a heterogeneity that has been previously noted^13,52,53^, but often ignored when reducing the analysis to a discrete outcome of pass and fail. Nica et al.^13^ noted that endpoint features and patterns derived from kinematic measures present a complex relationship in the healthy and post-injury stages, suggesting the use of distinct strategies to achieve the same endpoint performance by individual subjects. The authors called for the need for individualized methods of monitoring performance. In that sense, our subject-specific referencing, and the possibility to study KD-trajectory patterns per individual, provides a potential answer to that call. Our method assumes that the expected successful trial of a task while healthy is the best an individual is expected to perform. This assumption is reasonable in our experiments, and similar approaches, where subjects are trained on a task until they reach a plateau in performance. However, this assumption will not be true for untrained animals or for those that fail to train; however, KDI can also be used to understand how task acquisition develops with endpoint expert level performance used as a reference to evaluate early-stage performance. This subject-specific referencing is also an advantage for future use of our metrics for more naturalistic behaviors such as walking or food handling, where nuances in the heterogeneity of movement might be greater than in trained tasks. On the other side, subject-specific referencing is not applicable in situations where healthy performance is not available for the same individual, as often happens in clinical settings. One alternative could be to create references for a matched healthy cohort to our cohort of study, as it is done for many other metrics for the evaluation of humans. Future studies will need to be conducted to understand whether matched healthy individuals are a valid reference. Novel deep learning approaches that learn multidimensional representations of complex data conditioned to a set of desired covariates^54^ could be a promising approach to consider subject-specific characteristics in the analysis of movement.

One limitation of data-driven representation learning approaches, such as the one we used, is the difficulty in interpretating meaning from the low-dimensional features. We did not seek to explain the effects of injury in the low-dimensional space as that was out of our immediate scope, and because it is secondary to the value of KDI as a subject-specific differential metric of movement performance. Further work is needed to understand the relationship between the learned representations, the KD trajectories, and the sequence of movements that drives such representation. This could provide information on the different patterns or motor strategies that emerge during recovery after a functional disruption and be used as a tool to elucidate recovery and compensatory mechanisms.

Another important limitation is the normalization of time to a standard length. This was done for convenience during the development of the KDI, as differences in the trial completion time require further consideration. It is possible to consider the individual time of trajectory per trial in our framework as the reference, and the loadings of the PCA describe the per-trial low-dimensional space. It is also possible to extract those values independently of the number of frames per trial. This will be important in conditions that result in movement speed changes, such as the slowed movements observed in cerebellar injuries ^55,56^. However, the impact of the duration of the trial on the definition and interpretation of the KDI as a contrast to the healthy reference is unclear. Future work could explore improvements in our KDI algorithm to consider time in its true form. In addition, it is technically possible to combine the frame-by-frame location with derived metrics of interest, such as digit spread and paw angle per frame, frame-to-frame velocity and acceleration, and marker-to-marker distance, amongst others. We did not add those into our initial derivation of the KDI as it makes the interpretation of the metric considerably more difficult, and such further extensions should be paired with work on interpreting the learned representations.

## Acknowledgements

We thank Paola Di Grazia, MS, MHC-LP for technical assistance.

## Supplementary Figures

**Supplementary Figure 1.**
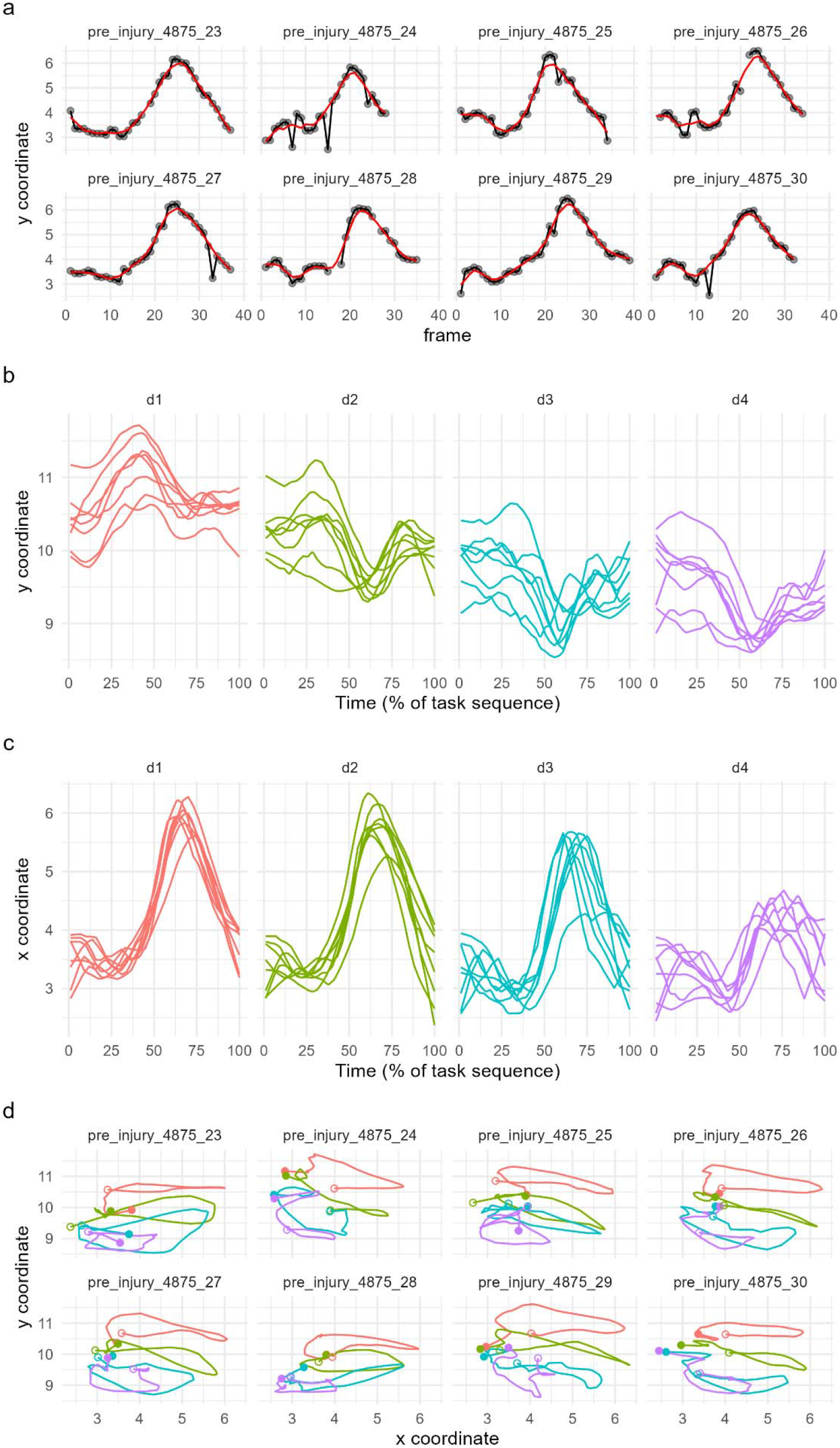
Example of the processing of the raw kinematic data for a set of trials from the same animal. (a) illustration of the denoising and smoothing of the y coordinate of the marker for d1 marker for the successful trials of the same animal from the first cohort pre-injury. The dots and black lines represent the original raw trajectory over the frame sequence, and the red line is the result of denoising and smoothing to consider spatiotemporal coherence. (b) The y coordinate of the time-normalized sequence for all four-digit markers of the same animal. (c) The x coordinate of the time-normalized sequence for all four-digit markers for the same animal. (d) The xy coordinate trajectories for the four-digit markers for the same animal. Close dots show the start of the sequence, and open dots show the end. Colors denote the different digits.

